# Near-cognate tRNAs dominate codon decoding times in simulated ribosomes

**DOI:** 10.1101/2025.02.04.636423

**Authors:** Fabio Hedayioglu, Emma J. Mead, Sathishkumar Kurusamy, James E.D. Thaventhiran, Anne E. Willis, C. Mark Smales, Tobias von der Haar

## Abstract

The codon sequence of messenger RNAs affects ribosome dynamics, translational control, and transcript stability. Here we describe an advanced computational modelling tool and its application to studying the effect of different tRNA species on the codon decoding process. We show that simulated codon decoding times are sensitive to the abundance of near-cognate tRNA species as well as cognate species, an aspect of the decoding system that is not fully considered in other computational modelling studies. We demonstrate that codon decoding times predicted by models that accurately define near-cognate tRNAs and that are parameterised with high-quality tRNA abundance datasets are highly similar to ribosome dwell times determined using experimental ribosome footprinting data, thereby confirming both the importance of near-cognate tRNAs for the codon decoding process and the general accuracy of our modelling tools.

## Introduction

Computational models of codon decoding during protein synthesis are widely used to predict and study gene expression. By enabling the detailed interrogation of specific molecules and their reaction rates, such models have helped elucidate basic features of the protein synthesis machinery in different species ^1–4^ as well as features of specific translational control pathways ^5,6^, have contributed to the study of the evolution of the translational machinery ^7–10^, and have informed the design of efficient recombinant sequences in biotechnology and medicine ^11–14^.

During codon decoding tRNAs are randomly sampled from the available cellular tRNA pool. tRNA sampling may deviate from a perfectly mixed system where all pool members are sampled with equal probability, for example, recently used tRNA species may be selected more frequently than expected by chance ^15^. However, any deviations from perfectly mixed systems are likely small, and sampling of the tRNA pool is clearly necessary for ribosomes to select cognate tRNAs. The fate of a tRNA that enters the ribosomal A-site is determined by the base pairing pattern between its anticodon and the A-site codon. Depending on the strength of base-pairing, sampled tRNAs may be released from the A-site, or may be accepted and transfer of the nascent peptide onto the amino acid carried by the sampled tRNA may be initiated. If peptidyl transfer occurs, this is followed by translocation of the ribosome onto the next codon, where the sampling process recommences ^16,17^.

For each of the 61 sense codons of the canonical genetic code, the cellular tRNA pool can be divided into different tRNA classes depending on the strength of codon: anticodon base pairing. Strongly pairing tRNAs that can initiate peptidyl transfer are designated as “cognates”, in contrast to “non-cognates” which base-pair less strongly or not at all and therefore do not initiate tRNA transfer. Moreover, available data suggest that tRNAs exist in more complex classes than the simple cognate/ non-cognate division. For example, some cognates exclusively form standard Watson:Crick base pairs whereas others form non-standard “wobble” base-pairs. Wobble pairing tRNAs can initiate peptidyl transfer, but in at least some cases do so with lower probability than Watson:Crick pairing tRNAs, and wobble-pairing cognate tRNAs can be rejected from the A-site ^18^. Similarly, anticodons of some non-cognate tRNAs do not base pair with the codon at all, whereas others base-pair in one or more of the three nucleotide positions. Where such partial contacts are strong enough to produce non-zero probabilities of initiating peptidyl transfer, the corresponding tRNAs are designated as “near-cognates” ^19,20^. While many studies have addressed the biochemistry and function of cognate tRNAs, both the nature of near-cognates and their role in shaping codon decoding and the genetic code is incompletely understood. For accurate protein synthesis the reliable acceptance of cognates and rejection of near- and non-cognates are equally important ^21^, and the relative lack of understanding of near-cognate tRNAs is thus an important knowledge gap.

Codon decoding models need to classify all possible anticodon: codon pairs in terms of the base-pairing patterns described above. Published approaches differ in which classes are considered as explicitly modelled species, with some models only considering the abundance of cognate tRNAs, whereas others consider additional species ^22,23^. Moreover, published modelling approaches also differ in the level of detail with which the codon decoding process is represented. Decoding of individual codons may be modelled as a step with a single rate parameter ^24,25^, as two separate codon decoding and translocation steps ^2,26^, or as more detailed processes where tRNA accommodation, GTP hydrolysis, peptidyl transfer and other sub-reactions occur with their own, separate rate parameters ^22,23^. The level of detail with which individual studies represent both tRNA classes and biochemical rate information depends on the study purpose and on the data sources used for parameterisation.

In the current study we extend the general principles we have used previously ^1,23^, to develop new modelling software that represents both tRNA classes and biochemical rate information at the highest level of detail. Moreover, these models can be parameterised flexibly, allowing users to explore previously inaccessible parameter spaces such as position-dependent changes in reaction parameters. We demonstrate the capabilities of the flexible parameterisation approach by exploring in depth how the representation of near cognates as a separate, difficult-to-reject class of tRNAs affects predicted codon decoding times.

Finally, we demonstrate that models based on accurate parameters for near-cognate tRNAs result in predicted codon decoding times that are highly similar to experimental ribosome dwell times extracted from ribosome footprinting data.

## Methods

### Model Structure

The new model developed and described herein uses fundamental reaction schemes which we and others have employed previously ^1,12,22^, and considers the four different tRNA classes outlined in the introduction (Watson:Crick pairing cognates, wobble pairing cognates, near-cognates and non-cognates). The consideration of two separate cognate classes is an extension of our previous model structure. The complete reaction scheme underpinning the model is summarised below (Figure 1).

**Figure 1.**
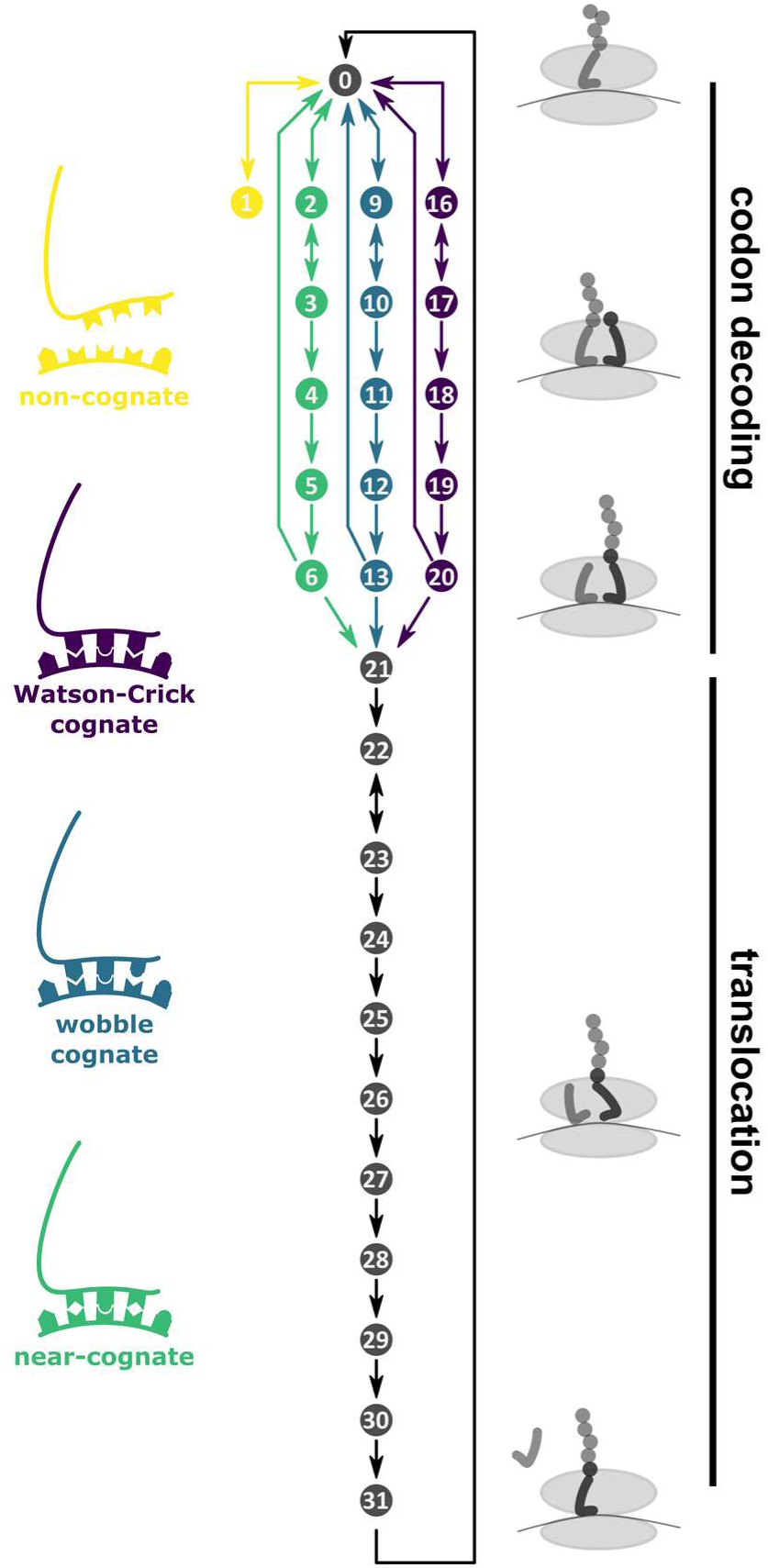
Reactions of the codon decoding cycle. Ribosomes with an empty A-site (state 0) can interact with four distinct classes of tRNAs, where the division of the total cellular tRNA pool into the four classes depends on the nature of the codon located in the A-site. Each ribosome undergoes transient interactions with tRNAs from the pool until aminoacyl-transfer has occurred (state 21), before the eEF2-catalysed translocation process begins which returns the ribosome to state 0 on the subsequent codon.

Individual tRNAs interact with the ribosome in a sequence of 16 consecutive reaction states that describe the outcomes of the various conformational changes, NTP hydrolysis, aminoacyl-transfer and translocation reactions that occur during a decoding cycle. The first six states comprise a series of reactions we term codon decoding which culminate in the aminoacyl transfer reaction, whereas the subsequent 11 states are termed translocation and include ejection of the P-site tRNA, transfer of the A-site tRNA to the P-site and forward movement of the ribosome by one codon.

Cognate and near-cognate tRNAs can undergo all of the ribosomal reactions, albeit with different rate constants which lead to different reaction rates and in consequence different propensities to reach the aminoacyl-transfer stage (Figure 1). To allow the modelling software to simulate these different rates efficiently, we define the reactions from initial tRNA binding to peptidyl transfer three times, once for each of the three corresponding tRNA classes. Thus states 2-6 correspond to the codon decoding phase for Watson-Crick decoding cognates, 7-13 to the codon decoding phase for wobble decoding cognates, and 14-18 to the codon decoding phase for near-cognates. Non-cognates never progress beyond the initial encounter reaction that leads to state 1. For any tRNA which reaches the amino acyl transfer stage rates are then independent of the class of the decoding tRNA, and we only assign a single series of state names to the corresponding reactions (states 21-31).

Since approximations for the rate constants with which the transitions between states occur are known, we can model this reaction scheme using standard rate constant based approaches including deterministic modelling based on differential equations, or stochastic modelling based on reaction propensities as proposed by Gillespie ^27^.

### Modelling Algorithms

We implemented two separate modelling algorithms, one dedicated to modelling the decoding of individual codons, and a second dedicated to modelling the decoding of mRNA sequences. Both were implemented as C++ software and are accessible via the Python classes CodonSimulator and SequenceSimulator. The CodonSimulator class is essentially an implementation of the Gillespie algorithm ^27^, and allows running stochastic simulations of the decoding of individual codons efficiently.

The SequenceSimulator class relies on a modified implementation of the Gillespie algorithm. A technical issue encountered in modelling the translation of mRNAs containing multiple ribosomes is how to represent the many different reactions that occur during the decoding of a transcript. The decoding of every single codon occurs via the reaction scheme outlined in Figure 1, so that the total number of distinct reactions related to a sequence of length *N* is *N\**31 (the number of reactions that can occur on each codon). Moreover, because standard Gillespie algorithms cannot track positional information, this set of reactions needs to be replicated for transcripts decoded by a single ribosome (monosomes), transcripts decoded by two ribosomes simultaneously (disomes), and all N-some variants that can potentially occur on a transcript. Using this approach, it was necessary to implement a system of 968 reactions and 242 components to model a three-codon transcript ^3^. While we and others have proposed agent-based approaches that circumvent the need for overly large reaction systems, here we systematically revised the implementation of the algorithm to improve computational performance.

The SequenceSimulator class works with three vectors. One vector contains the codon sequence of the transcript. A second vector contains the current state of each codon of the sequence, which can be either “<u>A</u>vailable”, “<u>U</u>navailable” i.e. covered by a ribosome although not located in the ribosomal A-site, or “<u>D</u>ecoding” i.e. located in a ribosomal A-site. A third vector contains simulator objects: a simulator object represents an individual ribosome, which is computationally represented as a data structure that contains the current reaction state of the codon, and the reaction parameter set for the A-site codon occupied by that ribosome. There is one set for each codon in “Decoding” state on the sequence, or for each ribosome (an N-some is therefore represented by a simulator vector of length N).

The information in these three vectors is processed by a simulation engine, which cycles through the following series of steps: 1) All possible reactions in the system are selected, including the possible reactions occurring with elongating ribosomes, but also initiation reactions which add a new simulator objects (ribosomes), and termination reactions which remove ribosomes or simulator objects from the vector. Reactions that involve changes in the state vector (i.e. ribosome movement, initiation or termination) are only allowed if they do not conflict with the location of another ribosome, i.e. ribosomes cannot move onto codons already occupied by other ribosomes. 2) Once the list of possible reactions and their rates has been established, the next reaction and the time required for that reaction are established using the “next reaction” approach ^28^. 3) The state and simulator vectors are updated to reflect any changes occurring from the execution of the reaction selected in step 2. This cycle is repeated until a pre-defined stop condition is met, which can be related to elapsed simulation time, computation time, or to a particular number of ribosomal termination events in the modelled system.

Our implementation provides distinct computational advantages, including that the parameters that need to be held in memory at any one time are limited, as well as operating advantages, for example that parameters can be very flexibly configured (this is explored in more detail in the Results section).

We further optimised the efficiency of the modelling software through additional measures. Due to the nature of the tRNA-dependent decoding system, the majority of tRNAs in the cellular pool have a “non-cognate” relationship with any given codon. Thus, the majority of tRNA: ribosome interactions are non-cognate binding events, and in fact if all non-cognate interactions are faithfully modelled, the majority of reactions occurring on a modelled transcript are binding and unbinding events of non-cognate tRNAs. Because non-cognates are processed with very rapid rate constants, these events only marginally contribute to the total simulated decoding time (this is explored in detail in the results section and in Figure 2). Thus, the software has a facility to remove non-cognate interactions from the scheme (non-cognate modelling is disabled by default but can be enabled by the user if desired).

**Figure 2.**
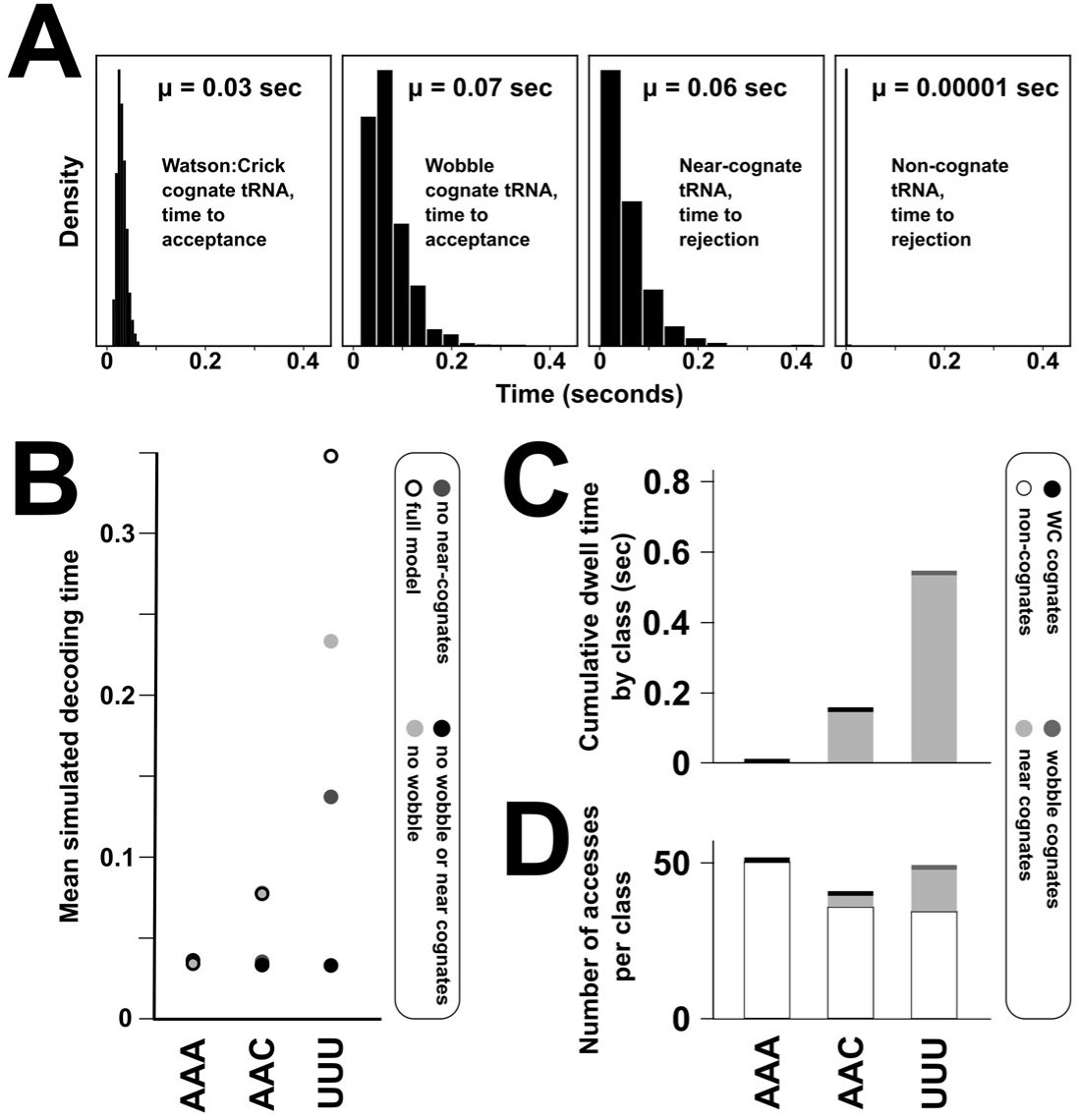
The effect of different model species on codon decoding times. **A,** processing times for individual tRNAs in the ribosomal A-site. **B,** predicted codon decoding times as a function of included species for three representative codons in the baker’s yeast decoding system. “Full model” refers to inclusion of separate Watson: Crick pairing cognate tRNAs, wobble-pairing cognate tRNAs, near-cognate tRNAs,and non-cognate tRNAs. **C,** the cumulative dwell time of each tRNA class during an average decoding process. Non-cognate dwell times are too short to be visible on the time scales of cognates and near-cognates. **D,** individual tRNA accesses per class in an average codon decoding cycle. For all codons non-cognates constitute the majority of sampled tRNAs, although these are rejected very rapidly and do not contribute strongly to the length of the decoding cycle.

As a second efficiency measure, we implemented a scheme that shortens the time required to simulate transcripts before they reach the steady state, for studies that are predominantly interested in answering questions about the steady state (as is the case for most published studies). If simulation runs are started with empty transcripts, simulations must run until the steady state is reached, and pre-steady state data are typically discarded. To start simulations much nearer to the steady state, we initially calculate the average decoding time for each individual codon, and pre-seed the modelled transcript with the ribosome density expected for a deterministic system. For example, if the initiation rate of the modelled system is 0.1 per second, we distribute ribosomes so that the codons between them take on average 10 seconds to decode. The resulting polyribosome complex is a system state that has the same ribosome density as the average stochastic state, provided that the effect of ribosome collisions in the system is small (collisions slow down ribosomes and increase ribosome density). The effect of this pre-seeding strategy is length-dependent, for a transcript of 500 codons modelled for 1,000 ribosome transits this reduces the required computation time by around 10-20% (Supplemental Figure 3).

### Model Parameterisation

Modelling the reaction scheme outlined in Figure 1 requires a number of parameterisation steps, including 1) establishing lists of known tRNAs to be included in the model as well as determining whether individual tRNA species belong to the cognate, near-cognate or non-cognate classes, 2) establishing abundance information for each tRNA species and converting this information into tRNA class abundances (the summed abundance in each of the two cognate, the near-cognate and the non-cognate tRNA classes for each codon), and 3) establishing rate information for each reaction in the scheme. A summary of the required input parameters is shown in Supplemental Figure 4. With a complete set of parameters, decoding times for individual codons can be modelled using the CodonSimulator class of the software. If the model is linked to a specific RNA sequence, ribosomal movement on the RNA can additionally be modelled via the SequenceSimulator class.

To provide a flexible tool that allows modelling of as wide a range of scenarios as possible, we provide a set of parameterisation tools that allow all model parameters to be freely configured. These are collected under the *set* methods of the simulator objects, including .*setPropensities(), .setInitiationRate(), .setTrnaConcentrations()* and others.

### Model Output

During simulations, the simulator software records all state changes with the times at which they occur in the modelled system. Recording every single reaction produces large amounts of data even though in many contexts it is predominantly the ribosome movement steps that are of interest, and the software therefore provides functionality to switch between coarse-grained state recording (change in ribosome position only) which preserves computing resources, and fine-grained state recording (all state changes in the system).

Both state change events and the time elapsed between them are recorded as variable vectors. In addition to the *event* and *time* vectors produced by both the codon and sequence simulators, the sequence simulator also records all ribosome collisions that occur during the simulation (all events where the distance between two ribosomes becomes zero). In addition, the primary simulation data can be used to calculate derived quantities through *get* methods, including ribosome transit times and individual codon decoding times for each codon.

### Software Application

The software is provided as a collection of Python wrappers that access and extend the function of the simulators which are written in C++. Detailed instructions for use are provided via Jupyter Notebooks that exemplify specific use cases and which are provided together with the analysis scripts and data accompanying this manuscript ^1^.

### Software Availability

For use on Windows and Linux systems the Python package(s) containing the modelling software can be downloaded free of charge and installed through the PyPI ^2^ distribution system. Source code is also freely available ^3^.

## Results

In addition to offering increased computational efficiency, our new ribosome simulators allow more flexible parameterisation compared to existing approaches. Here we demonstrate the capabilities of this flexible approach with an in-depth analysis of near-cognate tRNAs, a species that is not explicitly considered in many published approaches even though it strongly shapes the dynamics of codon decoding.

We initially explored the overall effects of inclusion or exclusion of different tRNA species, including near-cognates, on modelled codon decoding times (Figure 2). Individual tRNAs are typically processed by the ribosome within tens of milliseconds (Figure 2A), where “processing” means either undergoing all reactions up to and including peptidyl transfer (state 21 in Figure 1), or rejection from the A-site (a return to state 0). Only non-cognate tRNAs are released much more quickly than the other classes. The total decoding time for a codon is the sum of processing times for all tRNAs that need to be sampled before a tRNA reaches state 21 (the result of a peptidyl transfer reaction). Peptidyl transfer is then followed by ribosomal translocation, before the next tRNA sampling cycle begins.

When different tRNA species are included or excluded from simulations, predicted codon decoding times change for all codons to which these species are relevant. This is illustrated in Figure 2B for three codons that interact differently with the tRNA pool from baker’s yeast. In this organism AAA is decoded by the Watson:Crick pairing t(Lys)_UUU_ and according to the near-cognate definition we use, AAA does not have any near-cognate species (A is the nucleotide least able to undergo wobble interactions with unmodified nucleotides). AAC is decoded by the Watson:Crick pairing t(Asn)_GUU_, with t(Tyr)_GUA_ likely acting as a near-cognate. UUU is read by the wobble-pairing t(Phe)_GAA_, with multiple near-cognate species including (t(Ile)_AAU_, t(Ile)_GAU_, t(Leu)_GAG_ and t(Leu)_UAG_. In models based solely on the abundance of cognate tRNAs without distinction between Watson:Crick and wobble-pairing cognates, and where there is no distinction between near- and non-cognates, the decoding process is predicted to occur on very similar time scales for the three illustrative codons (Figure 2B, black points). If wobble-decoding tRNAs are modelled with their own specific rate constants ^29^, the decoding time for UUU (the only wobble-decoded codon in this analysis) is specifically reduced (Figure 2B, dark grey points). If near-cognates are modelled with their own rate constants ^30^, AAC and UUU codons are both predicted to be decoded more slowly (Figure 2B, light grey points), and distinguishing all four species leads to the greatest predicted differences in decoding times between the three codons (Figure 2B, open circles). Explicit inclusion of the different tRNA classes in models of the decoding process is thus necessary to capture the full dynamics of this process.

### Reaction rate sensitivity in codon decoding

To investigate how different tRNA species affect the dynamics of codon decoding in detail, we conducted a systematic sensitivity analysis for all model parameters. Starting with a basic parameter set as previously published ^12^, we varied each parameter within 10% of the original value and recorded how much codon decoding times for the 61 sense codons varied in response. This analysis yields an approximation of the formal control coefficients used in metabolic flux analyses ^31,32^.

Figure 3 illustrates how individual ribosomal reactions control codon decoding times, with positive correlation shown in blue, negative correlation in red, and grey indicating low or no correlation. For Watson-Crick pairing cognate tRNAs rate control resides predominantly in the initial formation of the tRNA:ribosome complex, consistent with the view that for strongly pairing cognate tRNAs the concentration (which is proportional to the rate of the initial encounter complex formation) drives the efficiency of codon decoding. For wobble-decoding tRNAs rate control is more strongly exerted at later reactions in the decoding scheme, in particular at reaction 3 and at the post GTP hydrolysis dissociation step. This is consistent with biochemical data which show that wobble decoding cognate tRNAs can be rejected at this step ^29^, in which case the entire sampling process must be repeated until the next cognate tRNA is encountered. Reducing rejection rates for wobble cognates by either increasing the rate of the peptidyl transfer reaction (reaction 6 in Figure 3) or by decreasing the rate of the dissociation reaction thus reduces the need to undergo the selection process repeatedly, and this controls codon decoding times more strongly than concentrations of wobble-pairing tRNAs. Lastly, for near-cognate tRNAs, those reactions that determine whether these tRNAs progress beyond the initial selection stages most strongly control codon decoding times (reactions 1-3 in Figure 3). Near cognate rejection is particularly slow if these species are rejected post GTP hydrolysis, and more frequent rejection at the initial stages speeds up the rejection process disproportionally.

**Figure 3.**
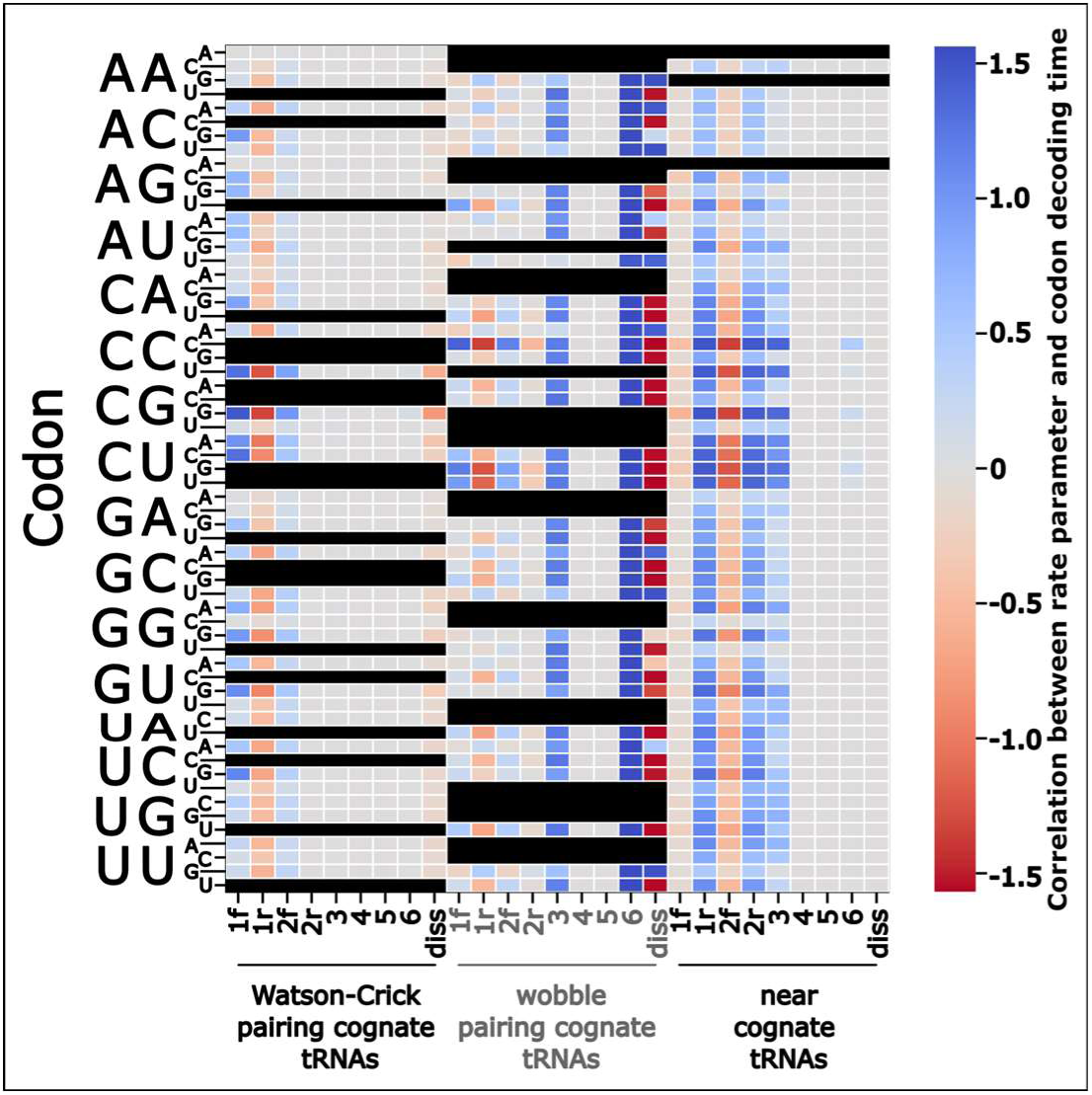
Sensitivity of codon decoding times on individual reaction rates. Colours indicate reaction rates that are correlated (blue), anticorrelated (red) or not correlated (grey) with simulated codon decoding times when individual reaction rates are altered within 10% windows.

In our analyses, some codons appear to show low or no sensitivity to any rate constants, most strongly those decoded only by a Watson-Crick pairing tRNA where no near-cognate species exist (codons AAA and AGA). For these codons a sampling cycle consists only of rejection of non-cognate tRNAs until the first cognate enters the A-site, which is almost always accepted. Because non-cognate rejection is very rapid this system behaves similar to a pure two-component system containing only translating ribosomes and cognate tRNAs, for which control coefficients associated with individual reactions are low (to significantly accelerate such a system one would have to accelerate *all* relevant rate constants simultaneously). Our results indicate that reactions exert strong rate control whenever they affect the ability of different tRNA species to compete, again highlighting that the full dynamics of codon decoding can only be understood if all relevant species are considered.

### Defining allowed base pairing patterns in near-cognate tRNAs

Plant *et al.* proposed a working definition for near-cognate tRNAs based on the ability of codon:anticodon pairs to form a Watson:Crick base-pair at the second codon nucleotide, in addition to either Watson:Crick or wobble base-pairs at the other two positions ^20^. Models defining near cognates in this way can rank sequences in order of expression levels with high accuracy ^12^.

Although models based on the Plant *et al.* definition are successful, there is remaining uncertainty regarding the kinds of wobble base-pairs that contribute to distinguishing near-cognate tRNAs from non-cognates. We explored the effect of different wobble base pairs on predicted codon decoding times, by systematically varying allowed base pairing contacts in our decoding model. Allowing or disallowing individual contacts affects which tRNAs are classed as near-cognates for a codon, thereby altering concentrations of its corresponding near-cognate tRNAs.

We initially compiled a comprehensive set of wobble base pairs known to form in solution between all modified bases occurring in yeast tRNA anticodons, and the four natural bases occurring in unmodified mRNA codons (Supplemental Figure 5). Two of these pairs, G:mcm^5^U and G:U, occur with essential wobble-decoding cognate tRNAs, and cannot be excluded from models without breaking the genetic code. The ten remaining base-pairs can be excluded while still resulting in a functioning genetic code, and we tested comprehensively how the inclusion or exclusion of these base pairs affected model performance. Simulating decoding times with all nucleotide contacts omitted in all possible combinations resulted in more than 700 datasets, each using differing definitions of near-cognate tRNAs *via* unique combinations of wobble interactions.

We initially investigated properties of this dataset by reducing its information content using principal component analysis (PCA) ^33^. Each dataset contains information on the decoding times for 61 sense codons. This high dimensionality can be reduced using PCA albeit at the cost of loss of information, for example, projecting information for the 61 codons into two principal components only allows recovering 61% of the available information (Figure 4A). Interestingly, the first seven components collectively capture more than 99% of the information of the full, 61-dimensional dataset, implying a subset of the 61 codons defines most of the information in the datasets. We hypothesize that this subset comprises codons that are particularly sensitive to interference from near-cognate tRNAs. We tested how much removal of any single codon from the datasets affected the amount of information recoverable in the first three principal components (containing 81% of the information of the full dataset). A small number of codons had significant effects on the analysis (Figure 4B), in that their removal increased the recoverable information. These analyses suggest that changing base pairing rules for near-cognates predominantly affects a subset of codons.

**Figure 4.**
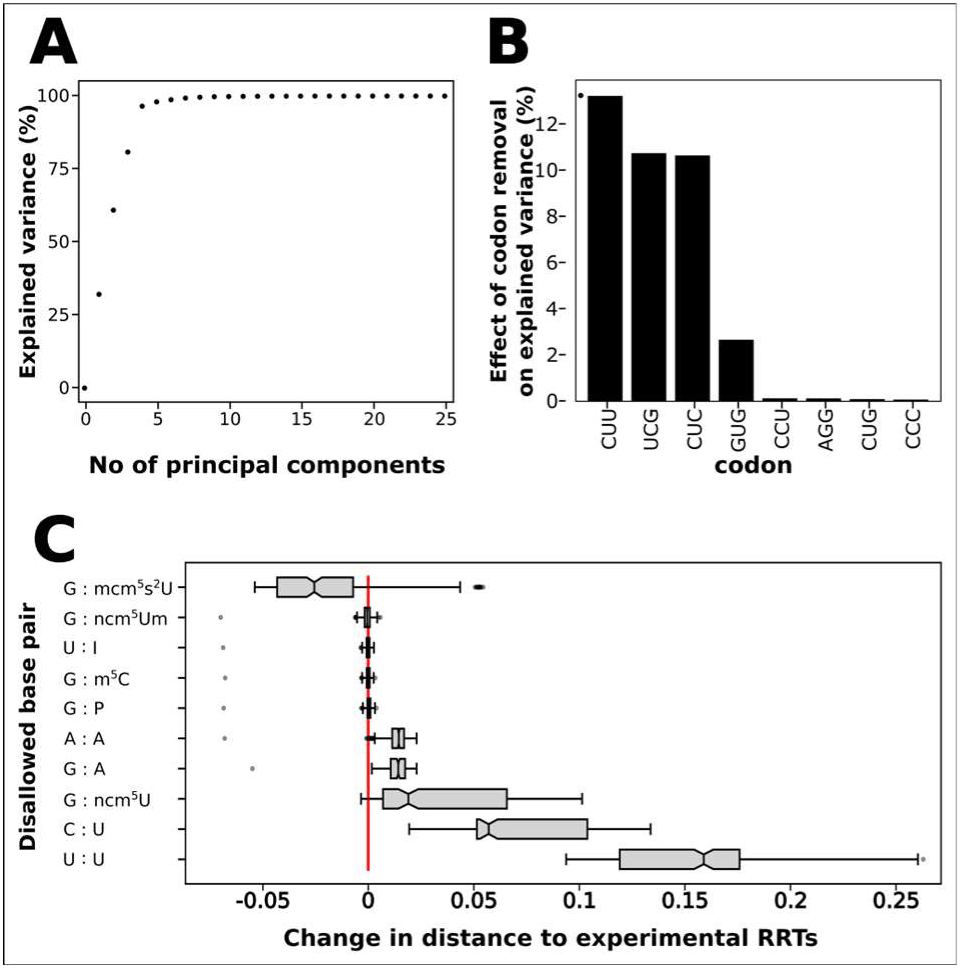
The effect of altered near-cognate definition on modelled codon decoding times. **A**, projection of the full, 61-dimensional datasets into reduced numbers of components leads to information loss for small numbers of components, but recovers >99% of information for more than six components. **B**, When individual codons are removed from the modelled datasets, the amount of information that can be recovered in the first three principal components increases for a small number of codons, indicating that these codons may be particularly sensitive to how near-cognate tRNAs are defined. **C**, removing individual wobble base pairs from the near cognate definition alters the similarity between predicted and experimentally determined codon decoding times.

We further explored relationships between datasets resulting from different base pairing rules by comparing predicted decoding times to experimentally determined “ribosome residence times” (RRTs). RRTs are estimates of the relative time decoding ribosomes spend on each codon derived from ribosome footprinting data, and have been reported by several groups for baker’s yeast ^34–38^.

Approaches for recovering RRTs from footprinting data are still debated, and data reported by different groups show variation (Supplemental Figure 6), but the degree of correlation between different datasets indicates that they contain at least some information on physiological codon decoding times. To compare these experimental datasets to our simulated decoding times we projected distances between the modelled and experimental datasets into 2-dimensional maps, initially exploring a variety of different approaches for dimension reduction and dataset normalisation (Supplemental Figure 7A). This exploratory analysis showed that multidimensional scaling (MDS, ^39^), when applied to mean normalised datasets, produced well separated clusters, with the number of clusters similar to the number of base pairs considered in the analysis, and with good correlation between mapped and original dataset distances (Supplemental Figure 7B). We therefore selected this approach for further analysis (Figure 5).

**Figure 5.**
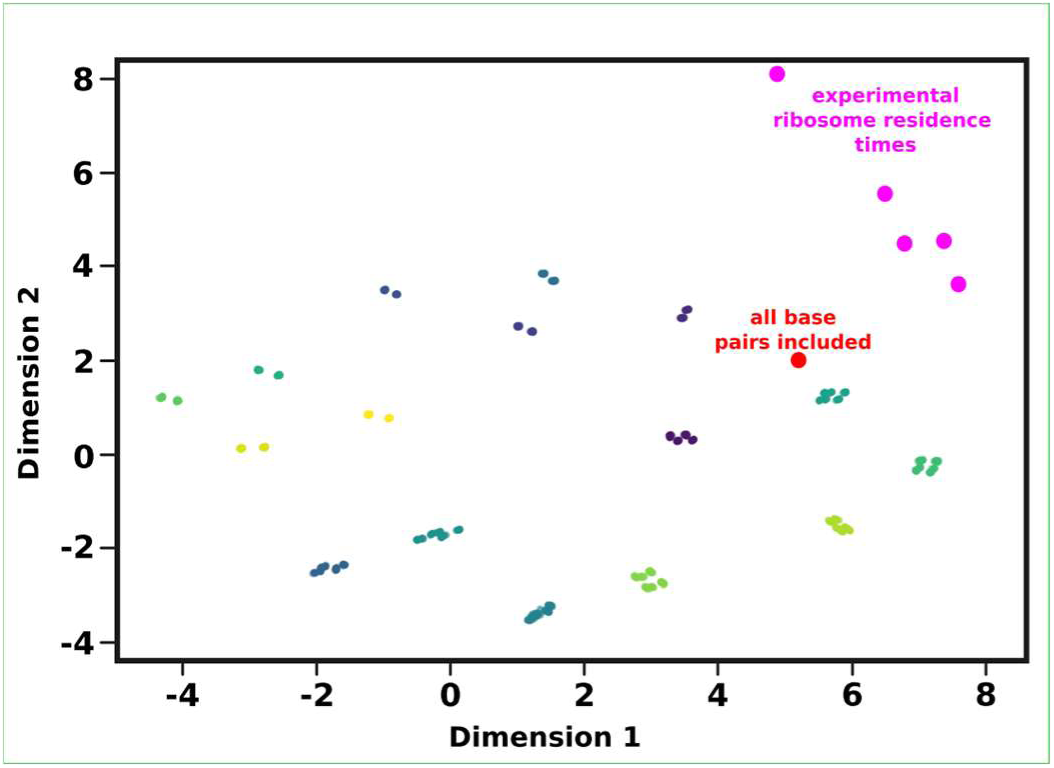
Comparison of codon decoding times modelled with different near-cognate definitions and experimentally determined codon decoding times. Experimentally determined codon decoding times are highlighted in magenta, modelled codon decoding times using a near-cognate definition that includes all possible wobble-base pairs is highlighted in red. All other data points are modelled codon decoding times where individual base pairs or combinations of base pairs were removed from the near-cognate definition, and pints are coloured to identify different clusters of similar modelling results.

In the map (Figure 5), different datasets are represented by individual dots, and distances between dots approximate the similarity between corresponding datasets. The modelled decoding times for the full set of base pairs and the experimental datasets are highlighted by red and magenta dots, respectively. The experimental RRT datasets show a spread in this mapping that relates to their quantitative differences. Codon decoding times predicted from simulations in which all candidate wobble base pairs are allowed to contribute to the definition of near-cognate tRNAs (red dot) are outside of the territory occupied by the experimental datasets, indicating that there are systematic differences between the model results and the experimental data, but the distance between the simulated data and four of the experimental datasets is smaller than the largest distance within the experimental datasets, indicating that the simulated dataset captures physiological codon decoding times with similar accuracy as individual experimental datasets.

When individual base pairs, or combinations of base pairs, are removed from the definition of near-cognate tRNAs, concentrations of near-cognates and in consequence modelled codon decoding times change (all points in Figure 5 other than the highlighted red and magenta points). None of these changes appear to reduce the distance between simulated and experimental data indicating that none of the changes make the simulated data more similar to the experimental datasets, and most changes increased the mapped distance between simulation results and the experimental datasets (Figure 5). At least some of the modelled base pairs are thus important to consider in the definition of near-cognate tRNAs, since their omission generates model predictions that become less similar to experimental results.

We validated these findings, and asked which codons produced the strongest effects on simulation results, by analysing what effect omission of individual base pairs had on Euclidean distances between the original, 61-dimensional datasets (Figure 4C). In this analysis the relative change in similarity to the experimental datapoints is quantified for omission of each wobble interaction, either in isolation or when paired with all possible combinations of other wobble interactions. Similar to the dimensional mapping (Figure 5), this analysis suggests that the predominant effect of removing individual base pairs from the near cognate definition is that simulation results become less similar to experimental results (i.e. the Euclidean distance to experimental results increases). Only the wobble interaction between guanine and 5-methoxycarbonylmethyl-2-thiouridine (mcm5s2U) produces a small net improvement in the similarity between simulations and experimental data, although this appears dependent on which other base pairs are also omitted. Omission of the uracil:uracil and cytosine:uracil wobble pairs affects the similarity between modelled and experimental data most strongly, indicating that these base pairs contribute most to the definition of near-cognate tRNAs, and inclusion or exclusion did not affect simulation results for four base pairs.

### Model-based evaluation of tRNA datasets

The quantification of cellular tRNA levels is challenging because the frequent occurrence of modified nucleotides interferes with the reverse transcription step required in most RNA quantification methods. Different approaches have been developed to negate the inhibitory effect of modified nucleotides, including alkaline hydrolysis which releases less modified fragments from full-length tRNAs (Hydro-tRNA-Seq ^40^), various strategies for generating full-length cDNAs despite the modifications (QuantMSeq ^41^, mimSeq ^42^, ARM-Seq^43^), and direct RNA sequencing by Nanopore ^44^. In baker’s yeast tRNA transcription is thought not to be strongly transcriptionally controlled, meaning that in this organism gene copy numbers constitute a useful approximation of tRNA levels ^45^, and all modelling data discussed so far were generated based on gene copy number counts.

A direct comparison of different tRNA datasets indicates that reported tRNA levels are strongly method-dependent (Figure 6A and Supplemental Figure 8). As a consequence, codon decoding times modelled based on the different tRNA levels also differ (Figure 6B). We mapped similarities between the different modelled codon decoding times and the experimental ribosome residence time datasets discussed earlier, using the same multi-dimensional scaling approach used above (Figure 6C). Gene copy number assessment, mimSeq and QuantMSeq methodologies predicted decoding times that were most similar to the experimental data, with Nanopore, ARMSeq and HydroSeq approaches producing modelled data that were less similar. Importantly, modelled data based on the gene copy number, mimSeq or QuantMSeq approaches map more closely to their nearest experimental dataset than the distance between the most divergent experimental datasets, suggesting that our computational model of the decoding process captures codon decoding times with similar accuracy as ribosome dwell time analyses based on ribosome footprinting data, provided they are parameterised using high quality information on tRNA abundance.

**Figure 6.**
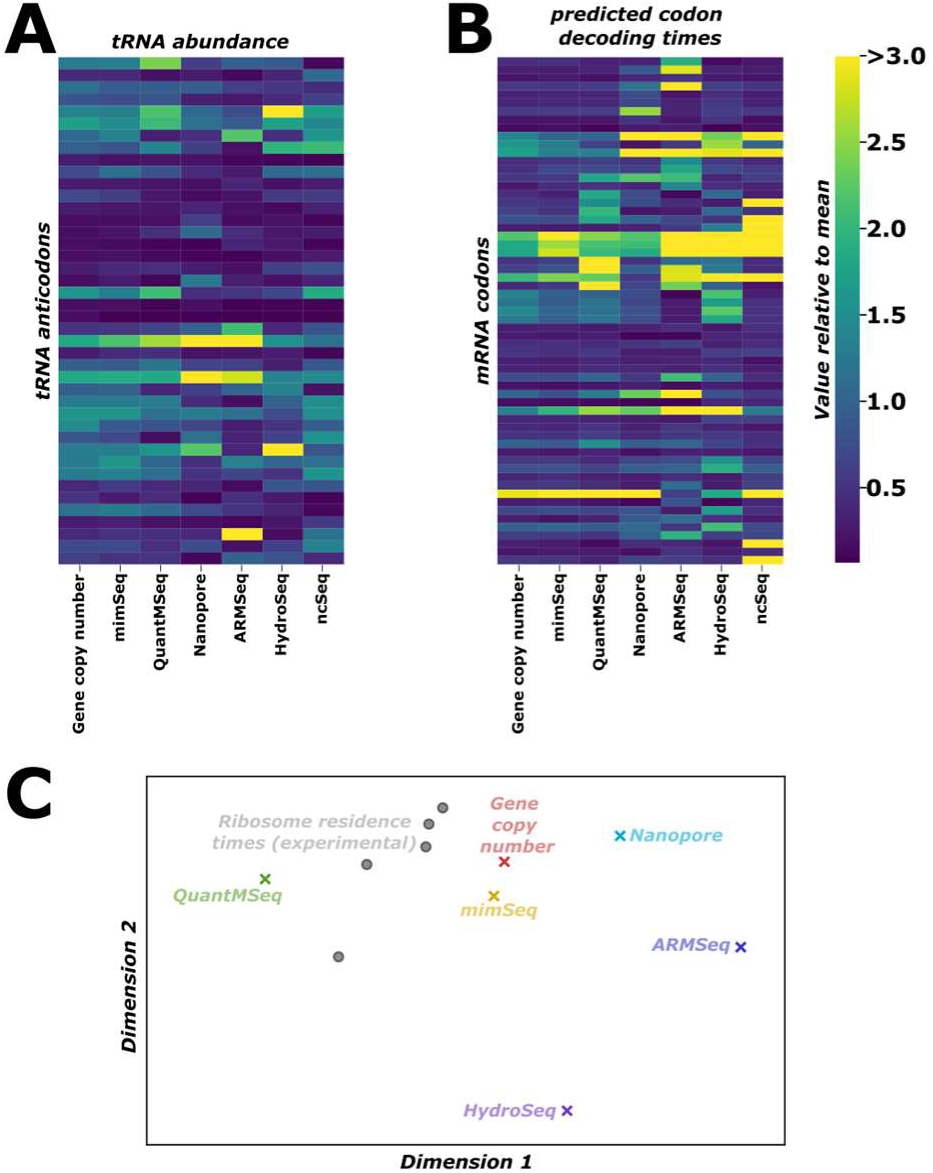
Comparison of model performance based on different tRNA datasets. **A**, tRNA abundances in datasets generated using different methodologies. **B**, codon decoding times simulated using the different tRNA datasets from A) as input. Data in A0 and B0 are normalised to sample means. **C**, 2-dimensional mapping of similarities between simulated codon decoding times (coloured markers) and experimentally determined ribosome residence times (grey markers). Distances between data scale with similarities between datasets.

## Discussion

In this study we demonstrate the capabilities of our new, flexibly parameterisable software for simulating ribosome- and tRNA-dependent codon decoding. The reliance on a principal model structure proven in other studies, together with the ability to freely adjust reaction parameters in an amino acid- and sequence-dependent manner, and with the performance advantages offered by an integrated modelling engine, enable easy modelling of scenarios that are not accessible with other, existing modelling software.

We demonstrate the capabilities of the software by exploring how near-cognates, a class of tRNA that has received less attention in past studies compared to cognates, fundamentally affect the dynamics of codon decoding. Introducing a near-cognate tRNA class that is rejected from ribosomal A-sites more slowly than standard non-cognates greatly reduces codon decoding times (Figure 2B), and for codons for which such tRNAs exist rejecting near-cognates from the ribosomal A-site typically becomes the most time-consuming process (Figure 2C). By how much the decoding process is slowed depends on the number of near-cognate tRNAs that need to be rejected on average before a cognate is accepted, which is determined by the near-cognate to cognate ratio.

Kothe and Rodnina showed that *E. coli* tRNA(Ala)_UGC_ nearly always progresses to the peptidyl transfer reaction once it reaches the late proof-reading step (the branch point at state 20 in Figure 1) on Watson-Crick pairing GCA codons, whereas the same tRNA is rejected 40% of the time on cognate but wobble-pairing GCC codons ^29^. Rejection of a cognate tRNA doubles the time required for codon decoding under the assumptions of a perfectly mixed system, since the entire sampling process (including slow rejection of any near-cognate tRNAs) needs to be repeated. Given the need to sample for extended times compared to Watson:Crick pairing cognates, high near-cognate:cognate ratios are likely particularly unfavourable for wobble-pairing cognates. This may explain why for amino acids where a single tRNA decodes two possible codons the genetic code shows some of the strongest biases against the wobble-decoded codon. A relevant example are GAA and GAG, both decoded by tRNA(Glu)_UUC_. The wobble-decoded GAG has two abundant near-cognate species, tRNA(Lys)_CUU_ and tRNA(Lys)_UUU_, and is one of most biased-against codons in the yeast genome with an average use ratio of the two glutamic acid inserting codons of 2.4:1^46^.

In addition to asking what overall effects near-cognate tRNAs have in the decoding process, we explored their definition in more detail. The data suggest that near-cognates are those tRNAs which can undergo partial base pairing between the codon and anticodon ^20,21,47^, and we adopted a working definition requiring formation of a Watson:Crick base pair between the central nucleotides, as well as wobble-base pairs at either or both of the other two positions ^20^. Which kinds of wobble-base pairs sufficiently stabilise contacts to make tRNAs behave like near-cognates is still unclear. To explore this question we chose a “brute force” approach in which we allowed or disallowed all possible wobble contacts that can be omitted without breaking the genetic code, in isolation or in combination with all other possible contacts. By comparing the resulting decoding time predictions to experimentally determined ribosome residence times from a number of published studies, we show that disallowing wobble contacts typically made predictions less similar to the experimental data (Figures 4 and 5), consistent with the notion that these contacts contribute to near-cognate behaviour in real tRNAs, and that such near-cognate tRNAs affect codon decoding times.

The salient parameter that distinguishes near-cognates from non-cognates (and equally wobble-decoding cognates from Watson-Crick decoding ones) is most likely the strength of base pairing between the tRNA anticodon and mRNA codon, which needs to be sufficiently strong to displace a series of “gatekeeper” nucleotides in the small subunit rRNA, thereby inducing conformational changes that are propagated throughout the ribosome ^20,48^. The probability of this displacement correlates with the affinity of the codon:anticodon pair, which explains both why weaker binding (wobbling) cognates can sometimes be wrongly rejected, while stronger binding near-cognates are more slowly and less reliably rejected than weaker non-cognates. Base-pairing strength is a continuous parameter, and we would expect that in reality individual tRNA:codon pairs are located on a continuum ranging from more near-cognate-like to more non-cognate-like, rather than behaving uniformly within two clearly separable classes. We currently lack the biochemical evidence to model this continuum in detail, but it is likely that the development of approaches that modulate the rate constants with which tRNAs interact in the ribosomal A-site based on the codon:anticodon affinity will further improve the accuracy with which codon decoding times can be predicted.

## Author contributions

EJM, SK, AEW, JEDT, CMS and TVDH designed the study. AEW, JEDT, CSM and TVDH obtained funding. FDLH wrote the modelling software. FDLH and TVDH conducted analyses and wrote the manuscript. All authors read and approved the manuscript.

## Acknowledgements

The authors gratefully acknowledge funding from the Wellcome Trust, UK (to AEW, CMS and TVDH, grant reference 201487/Z/16/Z); the Wellcome Leap R3 programme (to AEW, JEDT, CMS and TVDH), and the University of Kent Impact Acceleration Accounts (to TVDH, award references: Biotechnology and Biological Sciences Research Council (UK), BB/X511158/1, and Medical Research Council (UK), MR/X502753/1).

## Footnotes

1 https://github.com/tobiasvonderhaar/simulator_manuscript

2 https://pypi.org/project/elongation-simulator/

3 https://github.com/fheday/elongation_simulators

## Notes

https://github.com/tobiasvonderhaar/simulator_manuscript/

### Competing Interest Statement

The authors have declared no competing interest.

## References

1. Chu, D., Barnes, D. J. & von der Haar, T. The role of tRNA and ribosome competition in coupling the expression of different mRNAs in Saccharomyces cerevisiae. Nucleic Acids Research 39, 6705–6714 (2011).

2. Ciandrini, L., Stansfield, I. & Romano, M. C. Ribosome Traffic on mRNAs Maps to Gene Ontology: Genome-wide Quantification of Translation Initiation Rates and Polysome Size Regulation. PLoS Comput Biol 9, e1002866 (2013).

3. Matsuura, T., Tanimura, N., Hosoda, K., Yomo, T. & Shimizu, Y. Reaction dynamics analysis of a reconstituted *Escherichia coli* protein translation system by computational modeling. Proc Natl Acad Sci USA 114, E1336–E1344 (2017).

4. Rudorf, S. Efficiency of protein synthesis inhibition depends on tRNA and codon compositions. PLoS Comput Biol 15, e1006979 (2019).

5. Bastide, A. et al. RTN3 Is a Novel Cold-Induced Protein and Mediates Neuroprotective Effects of RBM3. Current Biol 27, 638–650 (2017).

6. Malik, Y. et al. Disruption of tRNA biogenesis enhances proteostatic resilience, improves later-life health, and promotes longevity. PLoS Biol 22, e3002853 (2024).

7. dos Reis, M. Solving the riddle of codon usage preferences: a test for translational selection. Nucleic Acids Res 32, 5036–5044 (2004).

8. Zur, H. & Tuller, T. Predictive biophysical modeling and understanding of the dynamics of mRNA translation and its evolution. Nucleic Acids Res 44, gkw764 (2016).

9. Deng, Y., de Lima Hedayioglu, F., Kalfon, J., Chu, D. & von der Haar, T. Hidden patterns of codon usage bias across kingdoms. J. R. Soc. Interface 17, 20190819 (2020).

10. Maheshwari, A. J., Calles, J., Waterton, S. K. & Endy, D. Engineering tRNA abundances for synthetic cellular systems. Nat Commun 14, 4594 (2023).

11. Racle, J., Overney, J. & Hatzimanikatis, V. A computational framework for the design of optimal protein synthesis. Biotech Bioeng 109, 2127–2133 (2012).

12. Chu, D. et al. Translation elongation can control translation initiation on eukaryotic mRNAs. EMBO J 33, 21–34 (2014).

13. Trösemeier, J.-H. et al. Optimizing the dynamics of protein expression. Sci Rep 9, 7511 (2019).

14. Ranaghan, M. J., Li, J. J., Laprise, D. M. & Garvie, C. W. Assessing optimal: inequalities in codon optimization algorithms. BMC Biol 19, 36 (2021).

15. Cannarozzi, G. et al. A Role for Codon Order in Translation Dynamics. Cell 141, 355–367 (2010).

16. Wilson, D. N. & Doudna Cate, J. H. The Structure and Function of the Eukaryotic Ribosome. Cold Spring Harbor Perspectives in Biology 4, a011536–a011536 (2012).

17. Knight, J. R. P. et al. Control of translation elongation in health and disease. Dis. Model. Mech. 13, dmm043208 (2020).

18. Agris, P. F., Vendeix, F. A. P. & Graham, W. D. tRNA’s Wobble Decoding of the Genome: 40 Years of Modification. J Mol Biol 366, 1–13 (2007).

19. Blanchet, S. et al. Deciphering the reading of the genetic code by near-cognate tRNA. Proc. Natl. Acad. Sci. U.S.A. 115, 3018–3023 (2018).

20. Plant, E. P. et al. Differentiating between Near- and Non-Cognate Codons in Saccharomyces cerevisiae. PLoS ONE 2, e517 (2007).

21. Manickam, N., Joshi, K., Bhatt, M. J. & Farabaugh, P. J. Effects of tRNA modification on translational accuracy depend on intrinsic codon–anticodon strength. Nucleic Acids Res 44, 1871–1881 (2016).

22. Fluitt, A., Pienaar, E. & Viljoen, H. Ribosome kinetics and aa-tRNA competition determine rate and fidelity of peptide synthesis. Comp Biol Chem 31, 335–346 (2007).

23. Chu, D., Zabet, N. & von der Haar, T. A novel and versatile computational tool to model translation. Bioinformatics 28, 292–293 (2012).

24. Dana, A. & Tuller, T. The effect of tRNA levels on decoding times of mRNA codons. Nucleic Acids Res 42, 9171–9181 (2014).

25. Shah, P., Ding, Y., Niemczyk, M., Kudla, G. & Plotkin, J. B. Rate-Limiting Steps in Yeast Protein Translation. Cell 153, 1589–1601 (2013).

26. Gritsenko, A. A., Hulsman, M., Reinders, M. J. T. & De Ridder, D. Unbiased Quantitative Models of Protein Translation Derived from Ribosome Profiling Data. PLoS Comput Biol 11, e1004336 (2015).

27. Gillespie, D. T. A general method for numerically simulating the stochastic time evolution of coupled chemical reactions. J Comp Physics 22, 403–434 (1976).

28. Gibson, M. A. & Bruck, J. Efficient Exact Stochastic Simulation of Chemical Systems with Many Species and Many Channels. J. Phys. Chem. A 104, 1876–1889 (2000).

29. Kothe, U. & Rodnina, M. V. Codon Reading by tRNAAla with Modified Uridine in the Wobble Position. Mol Cell 25, 167–174 (2007).

30. Mittelstaet, J., Konevega, A. L. & Rodnina, M. V. Distortion of tRNA upon Near-cognate Codon Recognition on the Ribosome. J Biol Chem 286, 8158–8164 (2011).

31. Kacser, H. & Burns, J. A. The control of flux. Symp Soc Exp Biol 27, 65–104 (1973).

32. Heinrich, R. & Rapoport, T. A. A Linear Steady-State Treatment of Enzymatic Chains. General Properties, Control and Effector Strength. Eur J Biochem 42, 89–95 (1974).

33. Jolliffe, I. T. & Cadima, J. Principal component analysis: a review and recent developments. Phil. Trans. R. Soc. A. 374, 20150202 (2016).

34. Gardin, J. et al. Measurement of average decoding rates of the 61 sense codons in vivo. eLife 3, e03735 (2014).

35. Pop, C. et al. Causal signals between codon bias, MRNA structure, and the efficiency of translation and elongation. Mol Syst Biol 10, 770 (2014).

36. Weinberg, D. E. et al. Improved Ribosome-Footprint and mRNA Measurements Provide Insights into Dynamics and Regulation of Yeast Translation. Cell Reports 14, 1787–1799 (2016).

37. Fang, H. et al. Scikit-ribo Enables Accurate Estimation and Robust Modeling of Translation Dynamics at Codon Resolution. Cell Systems 6, 180–191.e4 (2018).

38. Do Couto Bordignon, P. & Pechmann, S. Inferring translational heterogeneity from *Saccharomyces cerevisiae* ribosome profiling. FEBS J 288, 4541–4559 (2021).

39. Kruskal, J. B. Multidimensional scaling by optimizing goodness of fit to a nonmetric hypothesis. Psychometrika 29, 1–27 (1964).

40. Gogakos, T. et al. Characterizing Expression and Processing of Precursor and Mature Human tRNAs by Hydro-tRNAseq and PAR-CLIP. Cell Reports 20, 1463–1475 (2017).

41. Pinkard, O., McFarland, S., Sweet, T. & Coller, J. Quantitative tRNA-sequencing uncovers metazoan tissue-specific tRNA regulation. Nat Commun 11, 4104 (2020).

42. Behrens, A., Rodschinka, G. & Nedialkova, D. D. High-resolution quantitative profiling of tRNA abundance and modification status in eukaryotes by mim-tRNAseq. Mol Cell 81, 1802–1815.e7 (2021).

43. Cozen, A. E. et al. ARM-seq: AlkB-facilitated RNA methylation sequencing reveals a complex landscape of modified tRNA fragments. Nat Methods 12, 879–884 (2015).

44. Lucas, M. C. et al. Quantitative analysis of tRNA abundance and modifications by nanopore RNA sequencing. Nat Biotechnol 42, 72–86 (2024).

45. Ikemura, T. Correlation between the abundance of yeast transfer RNAs and the occurrence of the respective codons in protein genes. J Mol Biol 158, 573–597 (1982).

46. Nakamura, Y. Codon usage tabulated from international DNA sequence databases: status for the year 2000. Nucleic Acids Res 28, 292–292 (2000).

47. Kramer, E. B., Vallabhaneni, H., Mayer, L. M. & Farabaugh, P. J. A comprehensive analysis of translational missense errors in the yeast *Saccharomyces cerevisiae*. RNA 16, 1797–1808 (2010).

48. Ogle, J. M., Murphy, F. V., Tarry, M. J. & Ramakrishnan, V. Selection of tRNA by the Ribosome Requires a Transition from an Open to a Closed Form. Cell 111, 721–732 (2002).

