## Supplemental data for "Near-cognate tRNAs dominate codon decoding times in simulated ribosomes"

#### 1) Software performance

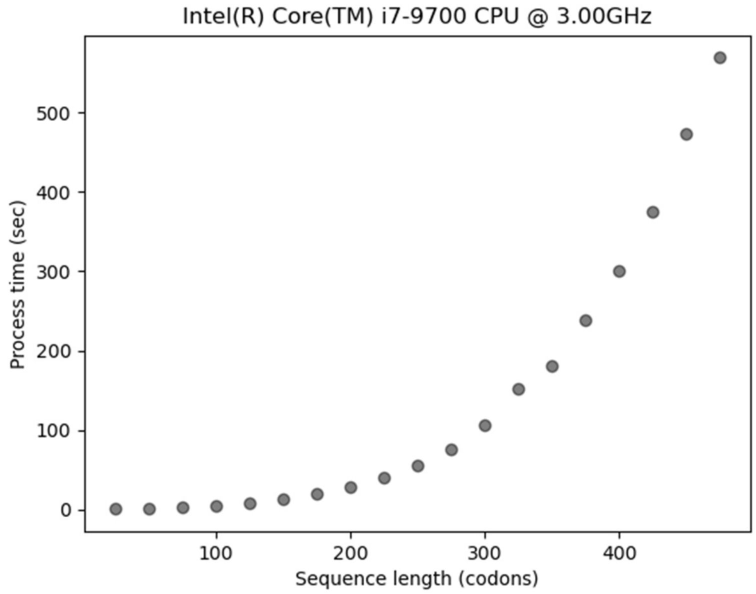


**Supplemental figure x. Modelling performance on a 1.8 GHz Intel i7 8565U CPU**. Sequences of increasing length encoding parts of Glutathione-S-Transferase were modelled using a termination criterion of 1000 terminated ribosomes.

#### 2) Inclusion of non-cognate tRNAs

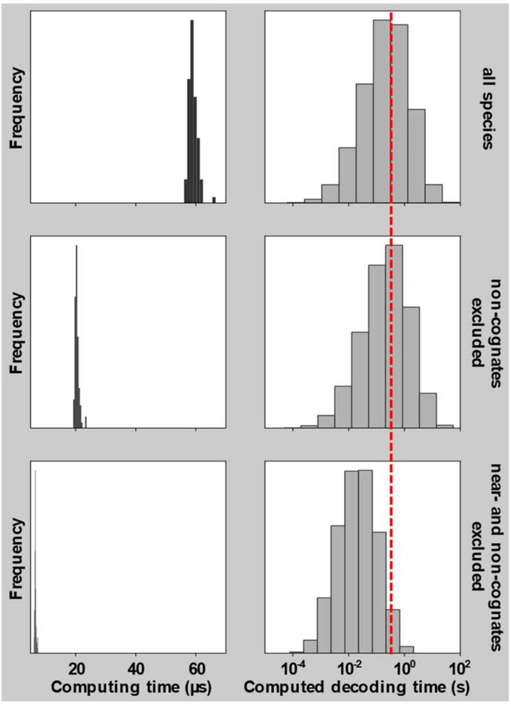


**Supplemental figure 2. Faithful mdoellign of all non-cognate interactions strongly increases computing time, without significantly affecting modelling results.** The top panel compares the computing time required (left) and the computed decoding time results (right) for a model that includes all species, including all non-cognate tRNAs. Omission of non-cognates reduces the computing time without affecting model results (center), whereas omission of near-cognates affects both computing time and model results (bottom).

### 3) Prepopulating transcripts

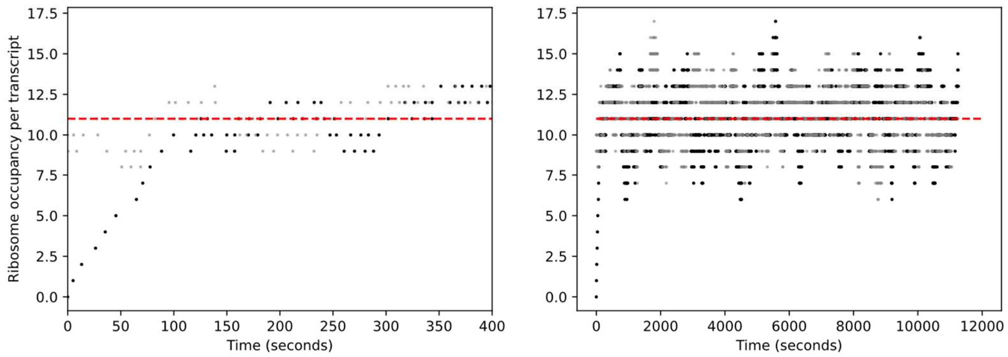


**Supplemental figure 3. Ribosome pre-seeding shortens pre-steady state simulation time.** Development of pre-steady state ribosome occupancies of model runs that begin with empty transcripts ( `sim.setPrepopulate(False)` , black dots), or with transcripts pre-seeded with a ribosome occupancy solution corresponding to the deterministic steady state solution ( `sim.setPrepopulate(True)` , grey dots). The panels show identical data from a simulation modelling translation of a yeast codon optimised glutathione-S-transferase gene, focusing on the first 400 seconds (left) or showing the full simulation (right). The red line shows the median ribosome occupancy from in the simulations excluding the first 250 seconds (11 ribosomes per transcripts in simulations both with and without ribosome pre-seeding).

### 4) Parameter requirements

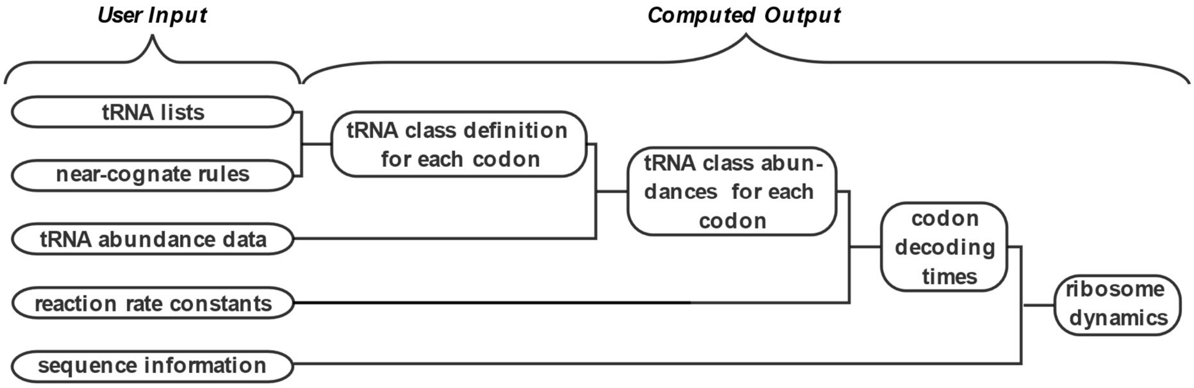


**Supplemental figure 4. Parameter requirements of the modelling software.**

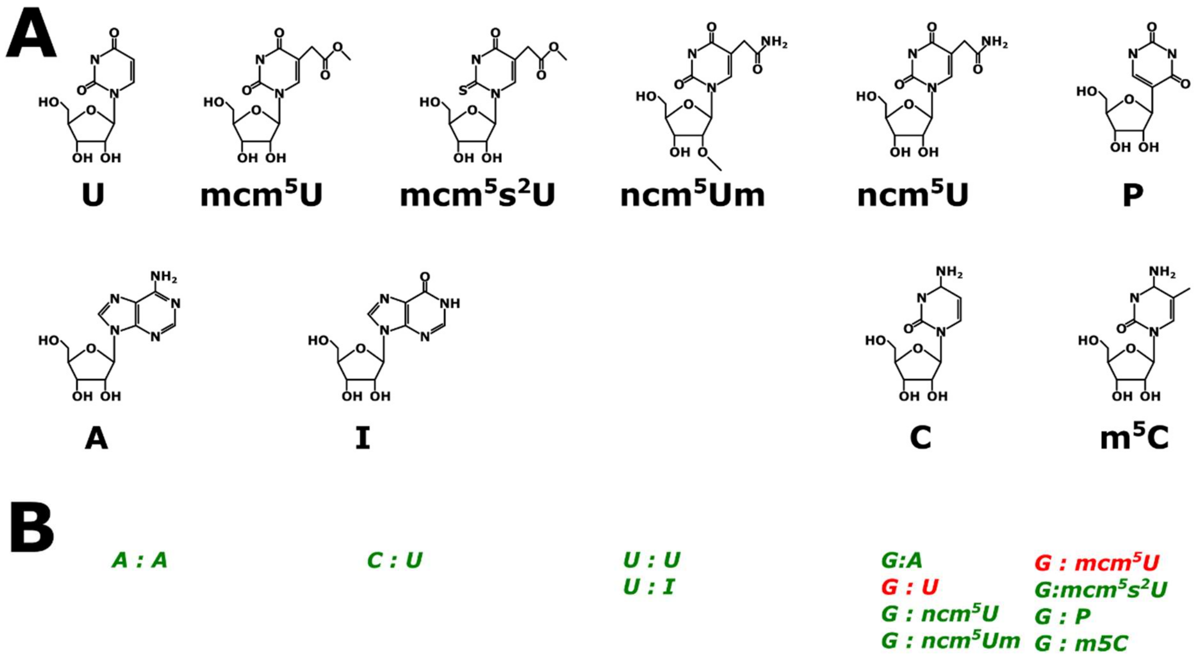


**Supplemental Figure 5. Modified nucleotides (A) and wobble base pairing interactions (B).** Colours in B) indicate base pairs that occur in essential wobble-pairing cognate tRNAs (red), and non-essential base pairs that were excluded in the analyses in figures 4 and 5 in the main manuscript.

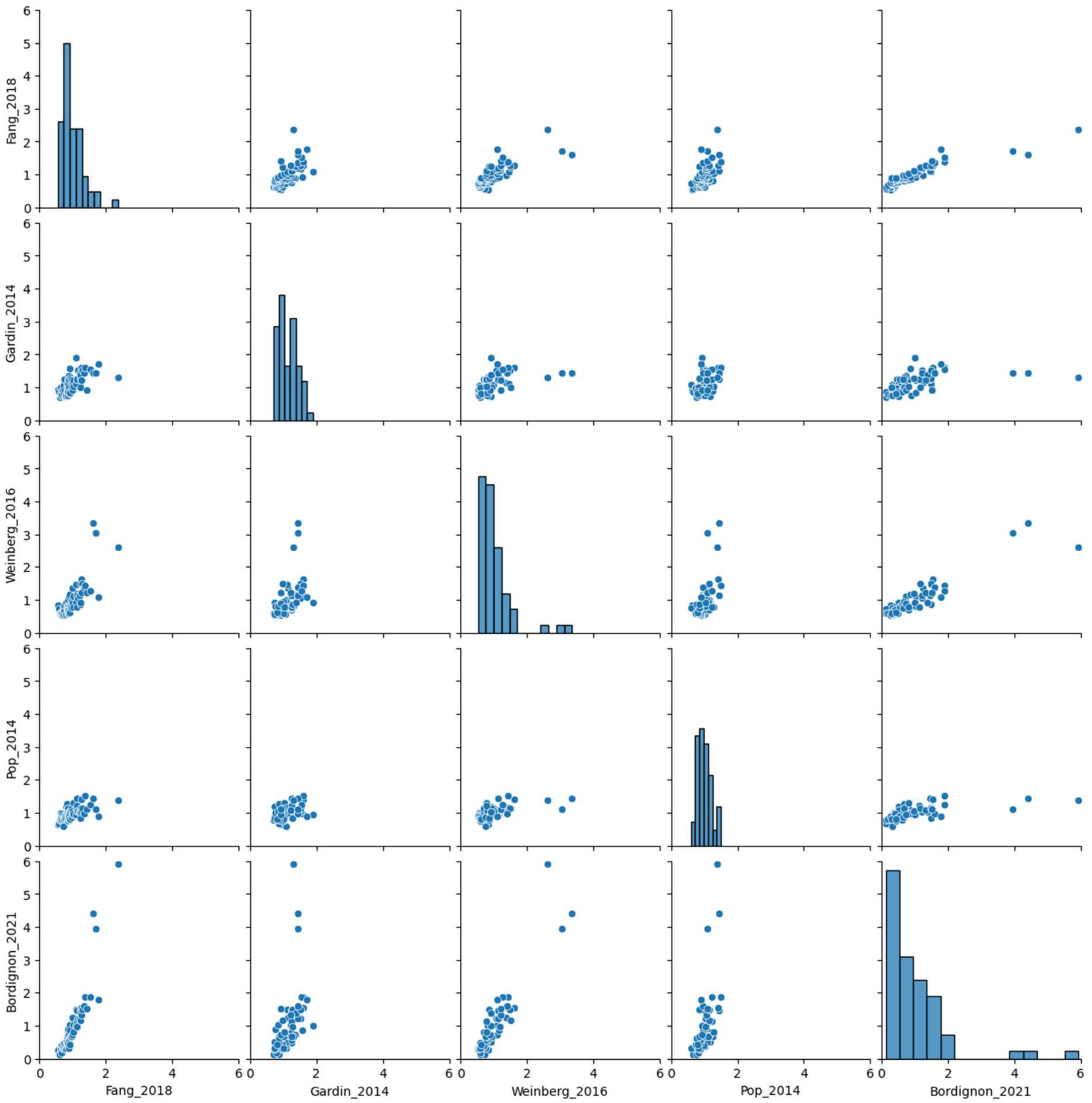


**Supplemental figure 6. Pairwise comparison of published Ribosome Residence Time (RRT) datasets included in this study.**

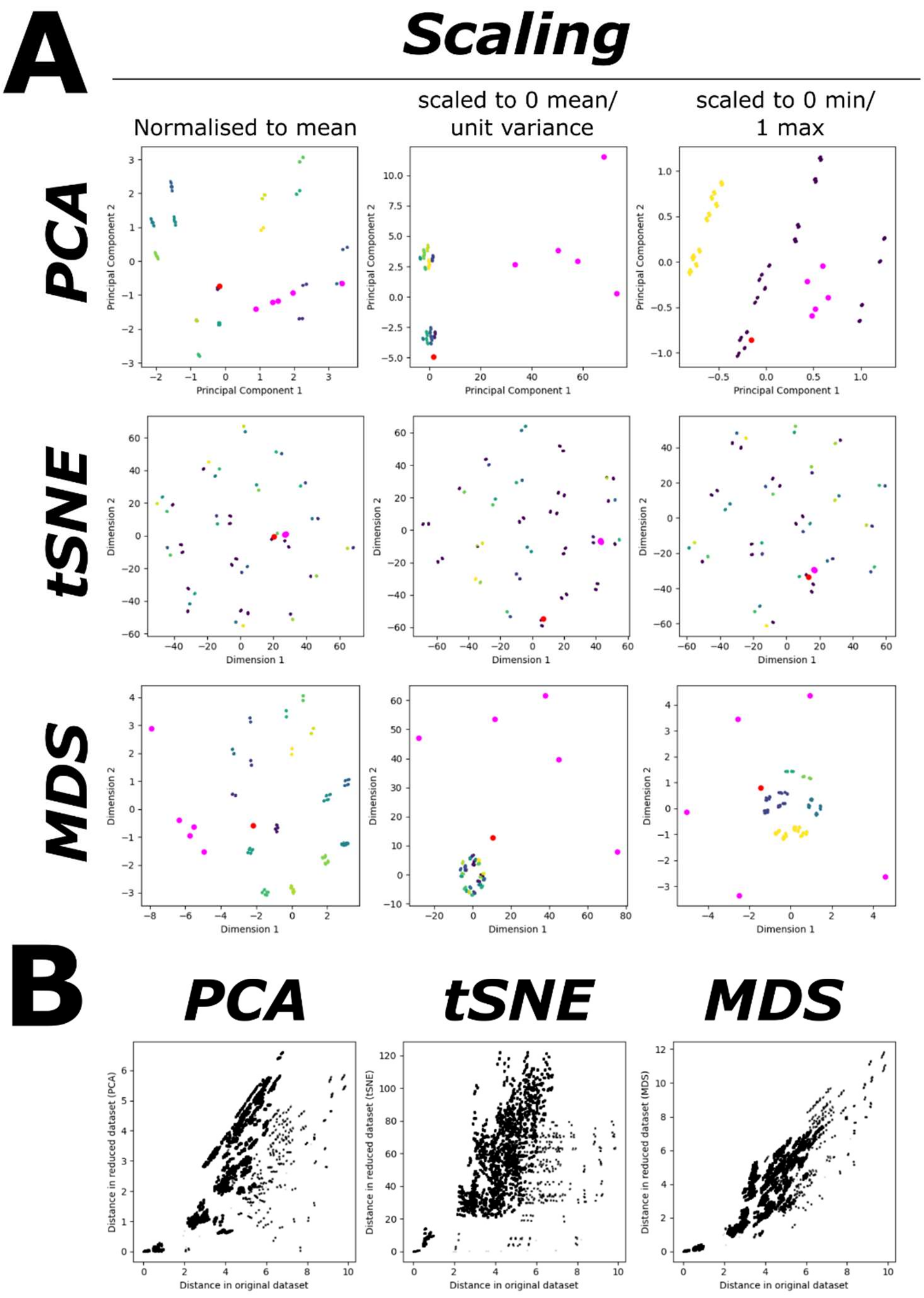


**Supplemental Figure 7. Dimension reduction approaches applied to modelled codon decoding times. A,** Principal Component Analysis (PCA), t-distributed Stochastic Neighbour Embedding (tSNE), and multi-dimensional scaling (MDS) were used to map Euclidean distances between modelled and experimental datasets, using different scaling approaches to normalise data. In all graphs, magenta data points represent experimental ribosome dwell time datasets; the red datapoint represents predicted codon decoding times with the full set of allowed nucleotide interactions, and all other datapoints represent modelled decoding times where one or more interactions were disallowed. Colours other than magenta and red identify datapoint clusters. Analyses in the main text refer to the MDS projection, for data which were normalised to the dataset mean. **B,** A comparison of Euclidean distances between the original datasets and between corresponding points in the different projections, for normalised to mean datasets.

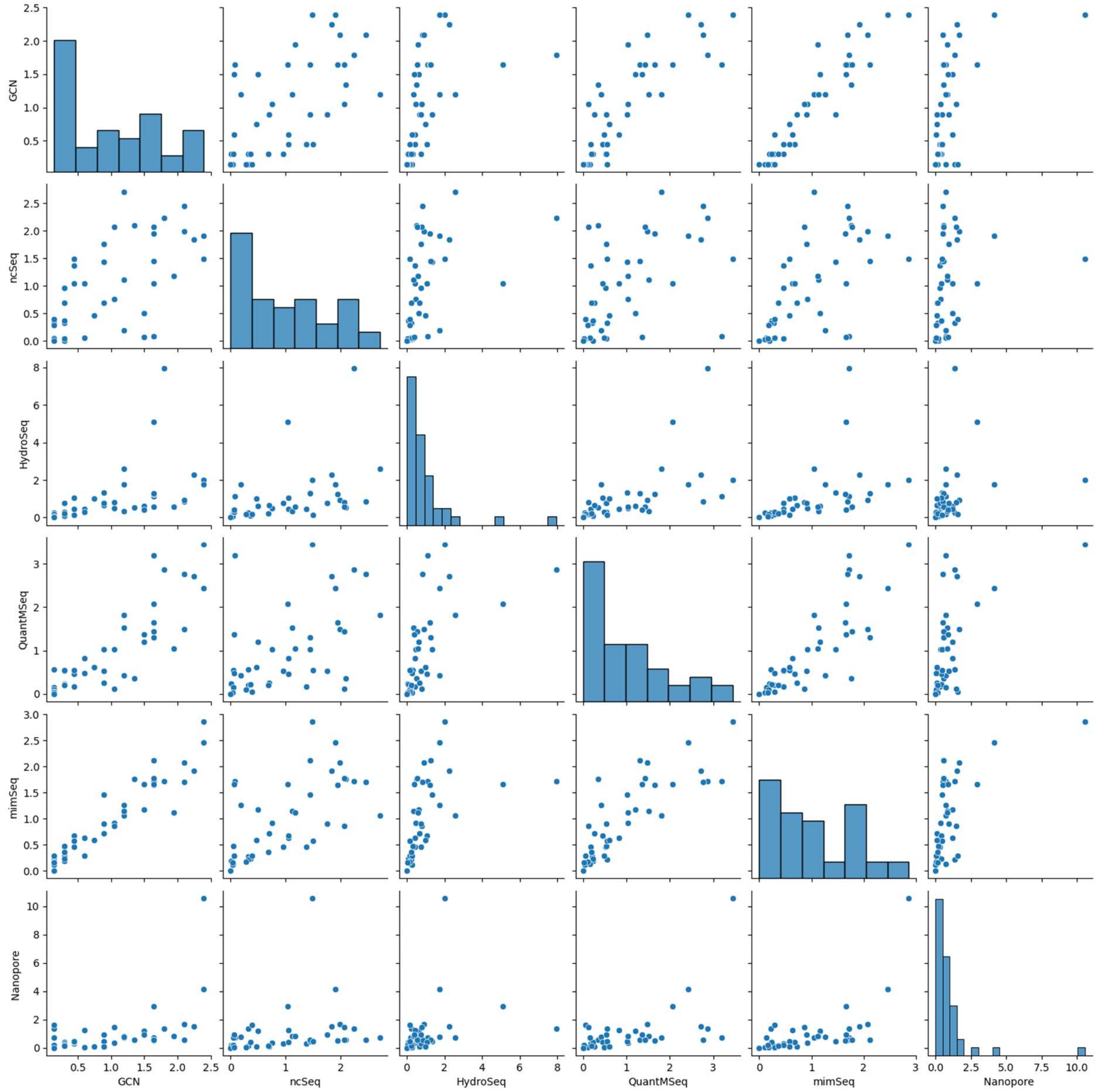


**Supplemental figure 8. Pairwise comparison of published yeast tRNA datasets.** Datasets are from NCBI GEO (ncSeq: xxx, HydroSeq: xxx, QuantMSeq: xxxx, mimSeq: xxxx), or from supplemental material of publications (Nanopore: Lucas *et al.* 2023). 'GCN' estimates tRNA levels based on the known gene copy number for each tRNA.
